# Domain-general enhancements of metacognitive ability through adaptive training

**DOI:** 10.1101/388058

**Authors:** Jason Carpenter, Maxine T. Sherman, Rogier A. Kievit, Anil K. Seth, Hakwan Lau, Stephen M. Fleming

## Abstract

The metacognitive ability to introspect about self-performance varies substantially across individuals. Given that effective monitoring of performance is deemed important for effective behavioural control, intervening to improve metacognition may have widespread benefits, for example in educational and clinical settings. However, it is unknown whether and how metacognition can be systematically improved through training independently of task performance, or whether metacognitive improvements generalize across different task domains. Across 8 sessions, here we provided feedback to two groups of participants in a perceptual discrimination task: an experimental group (N = 29) received feedback on their metacognitive judgments, while an active control group (N = 32) received feedback on their decision performance only. Relative to the control group, adaptive training led to increases in metacognitive calibration (as assessed by Brier scores) which generalized both to untrained stimuli and an untrained task (recognition memory). Leveraging signal detection modeling we found that metacognitive improvements were driven both by changes in metacognitive efficiency (meta-*d’/d’*) and confidence level, and that later increases in metacognitive efficiency were positively mediated by earlier shifts in confidence. Our results reveal a striking malleability of introspection and indicate the potential for a domain-general enhancement of metacognitive abilities.

## Introduction

Metacognition refers to the ability to monitor and introspect upon cognitive performance. An individual with good metacognition is aware of fluctuations in task performance, and appropriately modulates their confidence level (e.g. holding higher confidence when correct, and lower confidence when incorrect). While metacognitive abilities are often treated as stable characteristics of individuals (Allen et al., 2016; McCurdy et al., 2013; Fleming et al., 2010; Song et al., 2011), several lines of research hint at their malleability. For instance, practicing meditation boosts the accuracy of retrospective confidence judgments about recognition memory decisions (Baird et al., 2014) and monitoring of decision errors can be modulated by drugs (Hester et al., 2012) and brain stimulation (Harty et al., 2014). Moreover, recent work has identified distinct neural substrates in the frontal and parietal lobes supporting metacognitive monitoring across a range of tasks (Fleming et al., 2010; McCurdy et al., 2013; Baird et al., 2013; Allen et al., 2016; Cortese et al., 2017; see Fleming & Dolan, 2012, for a review), suggesting the potential for targeted modulation of metacognition independently of changes in first-order performance.

Previous attempts to improve metacognitive ability (confidence calibration) through explicit instruction, practice, feedback or a combination of these manipulations have led to mixed results, with some studies documenting increases, and others documenting null findings (e.g. Adams & Adams, 1958; Lichtenstein et al. 1982; Sharp et al. 1988; Bol et al., 2005; Nietfeld & Schraw, 2002; Renner & Renner, 2001). One potential explanation for such heterogeneity of results is that training may impact first-order performance, thus masking subtle changes in metacognition because they are positively correlated (Fleming & Lau, 2014; Sharp et al. 1988). Recent developments in the analysis of confidence-rating data now permit the effective isolation of metacognitive ability (the relationship between performance and confidence) from changes in performance through calculation of the signal detection theoretic parameter meta-*d’* (Maniscalco & Lau, 2012; Fleming & Lau, 2014). Because meta-*d’* is in the same units as first-order performance (*d’*) a metacognitive “efficiency” score (meta-*d’/d’*) is straightforward to calculate and indexes an individual’s metacognitive capacity with respect to a particular level of task performance. While training paradigms have proven effective in other cognitive domains, such as working memory (Klingberg, 2010; Morrison and Chein, 2011; von Bastian and Oberauer, 2014; Constantinidis and Klingberg, 2016) and even perceptual domains such as synaesthesia (Bor et al., 2014), it remains unknown whether metacognitive efficiency can be improved with practice, and whether putative metacognitive training supports transfer to untrained tasks or domains. Given that effective monitoring of performance is deemed important for effective behavioural control (Nelson & Narens, 1990; Metcalfe & Finn, 2008), intervening to improve metacognition may have widespread benefits, for example in educational and clinical settings.

However, it remains unclear whether such an intervention is *a priori* plausible for alleviating metacognitive deficits, or enhancing baseline metacognitive performance, across a range of scenarios. There is disagreement about the extent to which metacognitive ability is a domain-general resource that can be applied to multiple different tasks, or whether it is comprised of domain-specific components. Recent findings suggest that confidence is encoded in a “common currency” that can be compared across a range of arbitrary decision scenarios (de Gardelle & Mamassian, 2014; Faivre et al., 2017). However other studies indicate a substantial fraction of individual variation in metacognitive ability is domain-specific (Kelemen et al., 2000; Morales et al., 2018), consistent with dissociable neural correlates of perceptual and memory metacognition (McCurdy et al., 2013; Baird et al., 2013; Fleming et al., 2014; Morales et al., 2018). To the extent to which metacognition is domain-specific, training in one domain (for instance, on the computerized perceptual discrimination task that we employ here) may provide only narrow benefits to metacognition in that domain and be of limited value outside the laboratory. To evaluate the potential benefits of training on metacognitive ability it is therefore critical to assess whether such improvements generalize to an untrained task or cognitive domain. A useful parallel can be drawn with the literature on working memory training – here, meta-analysis suggests that “near” transfer to closely related tasks is commonly obtained, but evidence for far transfer is less consistent (Melby-Lervag & Hulme, 2013). The transfer profile of metacognitive training remains unknown.

Here we sought to investigate these questions by providing differential feedback to two groups of participants over eight training sessions on a perceptual discrimination task. A control group received feedback on their objective perceptual discrimination performance, whereas an experimental group received feedback on the calibration of their metacognitive judgments with respect to objective performance. Despite both groups exhibiting similar task performance, the experimental group displayed selective enhancements of metacognitive calibration (the association between confidence and performance) on the trained task. Furthermore, we obtained evidence for a transfer of metacognitive enhancements to an untrained stimulus type and to an untrained task (recognition memory). Together our results reveal a hitherto unreported malleability of domain-general mechanisms supporting metacognition and highlight the potential for generalized improvements in metacognitive ability.

## Methods

In this section we report how we determined our sample size, all data exclusions, all manipulations and all measures in the study (Simmons et al., 2012).

### Participants

We set out to recruit at least 30 subjects per group (60 in total), and no data were analysed prior to completion of data collection. Data were collected via Amazon Mechanical Turk (https://www.mturk.com), an online crowdsourcing platform. 102 adult participants completed at least the first session of the study. Of these, 8 participants were excluded from further training due to floor or ceiling performance in the pre-training baseline session, and a further 25 participants exited the study before completing the full training protocol. Of the remaining 69 participants, one was excluded due to technical problems and 7 were excluded based on data quality criteria explained in detail below. Final analyses were carried out on a dataset of 61 participants (35 women, 26 men, mean age = 38.1 years, age range: 20 – 64 years). Participants were required to use either Google Chrome or Mozilla Firefox in full-screen mode to complete the experiment on a computer(s) of their choosing.

Before participating in each session, all subjects provided informed consent as approved by the UCLA Institutional Review Board (IRB#15-001476). Subjects received monetary compensation in U.S. Dollars (*range* = $37.60 – $44.60) for approximately 5 hours (*mean* = 5.33 hours) of participation over a period of 9-35 days (Control group mean = 15.5 days, Experimental group mean = 15.4 days; independent samples t-test t(59) = 0.10, *p* = 0.92).

### Overview of procedure

The experiment was divided into 3 phases: Phase 1, pre-training (1 session) → Phase 2, training (8 sessions) → Phase 3, post-training (1 session), resulting in 10 sessions in total. Figure 1B provides an overview of the experiment timeline. Phase 1 consisted of stimulus titration and a pretraining session to evaluate baseline metacognitive accuracy in a series of 2-alternative forced-choice (2AFC) discrimination tasks (see *Task* below and Figure 1A). One set of tasks assessed perceptual discrimination, the other set assessed recognition memory. The tasks followed a 2×2 factorial design crossing cognitive domain (perception or memory) with stimulus type (explained in detail below).

**Figure 1.**
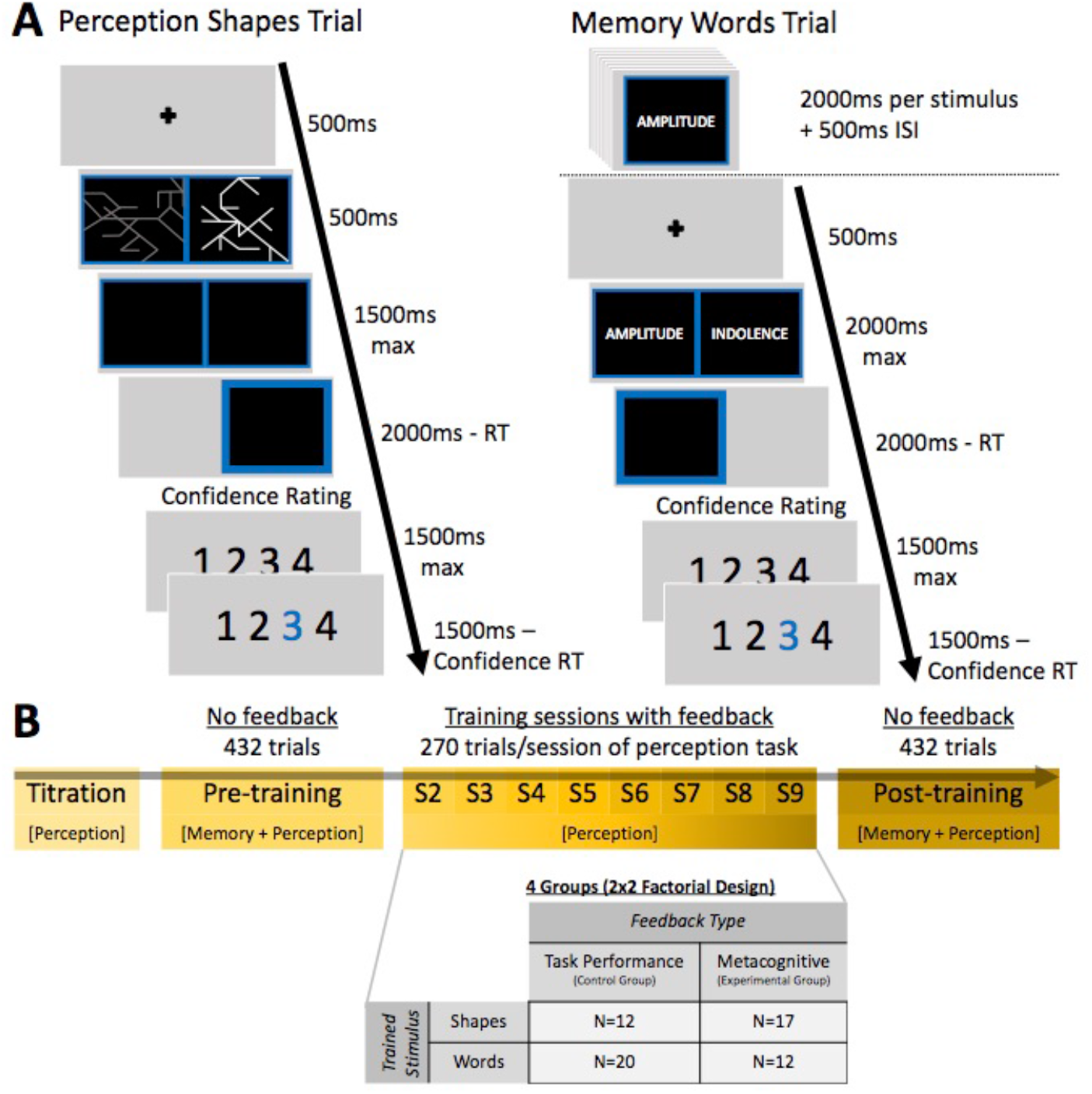
Task and session structure. **A)** Subjects were tested on both a perceptual discrimination and recognition memory task, each involving two stimulus types: abstract shapes and words. The perceptual task (left) comprised a 2-alternative forced-choice discrimination judgment as to the brighter of two simultaneously presented stimuli on each trial. The memory task (right) comprised an encoding phase followed by a series of 2-alternative forced-choice recognition memory judgments. **B)** Experiment timeline. Each subject completed 10 sessions in total: a pre-training session, 8 training sessions, and a post-training session. All four conditions were assessed at pre-and post-training, but only the perceptual task with a single stimulus type (shapes or words) was trained during sessions 2-9. During training sessions, the control groups received feedback on their objective perceptual discrimination performance, whereas the experimental groups received feedback on their metacognitive calibration. In both groups, feedback was delivered every 27 trials (see Methods).

Each task consisted of 108 trials, giving 432 total trials in the pre-training session. The order of these tasks was counterbalanced such that each participant performed both tasks in one domain followed by both tasks in the other domain, and within each domain the order of stimulus types was also counterbalanced.

At the start of Phase 2 subjects were assigned to one of four groups. Each group formed a cell in a 2×2 factorial design crossing feedback type (Control group vs. Experimental group) and trained stimulus type (see *Training Procedure* below). All subjects received training on the perceptual task only, with the recognition memory task introduced again at post-training to assess transfer to a different task domain. During the training phase, each of the eight sessions consisted of 270 trials (2160 trials total), and block-wise feedback was administered every 27 trials (see *Feedback* below).

Phase 3, the final post-training session, was identical to the pre-training session Phase 1 except that stimulus titration was omitted. Task order was counterbalanced against that used in pretraining, such that each subject performed the task domains (memory, perception) in the opposite order to that seen in pre-training. The order of stimulus types within each domain remained the same.

Phase 1 lasted approximately 60 minutes, the eight training sessions in Phase 2 lasted approximately 25 minutes each, and Phase 3 lasted approximately 45 minutes. Subjects were required to wait a minimum of 24 hours between each session and were asked via email to complete each subsequent session within 48-72 hours of the previous session.

### Tasks

Figure 1A displays example trial timelines for the perception and memory tasks. In the perception task, participants were presented with two images (either ‘words’ or ‘shapes’) and asked to judge “which [image] has brighter lines?” In the memory task, participants were first presented with a series of images to memorize (again either ‘words’ or ‘shapes’). On each subsequent trial, one old image and one novel image were presented with the instruction to judge “which [image] have you seen before”. In all tasks, after each decision, subjects were asked to rate their confidence on a 1-4 scale. They were informed that 1 corresponds to “very low confidence”, 2 to “low confidence”, 3 to “high confidence” and 4 to “very high confidence”.

In the pre-training session, before beginning each task, subjects completed 3 practice trials to become acquainted with making perception/memory judgments and using the confidence rating scale. Following the practice trials, we probed knowledge of how to perform the perception/memory judgments with a comprehension question asking “In the perception/memory task, how do you decide which image to choose?” The three response options were “which one you remember”, “which has more lines”, and “which is brighter”. If a participant answered either question incorrectly, they were excluded from further participation and offered a partial reimbursement determined by the proportion of the session completed. There were no practice trials or comprehension questions in the post-training session.

### Training Procedure

The second phase of the study involved eight training sessions of 270 trials each (2160 trials in total), spread over 8-34 days. Participants were randomly allocated to one of four groups in a 2×2 factorial design crossing feedback type (Control group vs. Experimental group) and trained stimulus type (shapes or words). All groups received block-wise feedback in the form of reward (points) every 27 trials. The Control groups (for both stimulus types) received feedback on their objective perceptual discrimination performance; the Experimental groups (for both stimulus types) received feedback on their metacognitive calibration, as determined by the average Quadratic Scoring Rule (QSR) score. The QSR provides a metric for how closely confidence ratings track accuracy (Staël von Holstein, 1970), and is equal to one minus the Brier score (Fleming & Lau, 2014). The rule underpinning each feedback type is described in more detail under *Feedback* below.

To ensure that each group fully understood how points could be earned, instructions were provided on the meaning of the feedback schedule. Participants completed eight demonstration trials which explained how earnings changed based on their objective performance (Control group) or the correspondence between confidence and accuracy (Experimental group). After the demonstration, subjects performed ten practice trials in which they received full feedback and a brief explanation. Note that in the demonstration and practice trials, feedback was calculated on a trial-by-trial basis and therefore differed from the block-wise feedback received in the training sessions (see *Feedback).*

After the demonstration and practice trials, participants were asked two comprehension questions probing their understanding of how to earn points. If they failed these questions they were asked to attempt them again until they were successful.

### Task Performance Titration

Throughout the entire 10-session experiment, the performance of each subject was titrated online to achieve approximately 75% correct for all tasks except the memory-words task. This “threshold” level of percentage correct produces sufficient trials for each Signal Detection Theory outcome (hits, misses, false alarms and correct rejections) for analysis of *d’* and meta-*d’* (Maniscalco and Lau, 2012), and ensured any changes in metacognitive sensitivity were not confounded by shifts in task performance.

Titration was accomplished in different ways for each task. In the perception tasks (for both word and shapes), we implemented two interleaved, weighted and transformed staircase procedures on the brightness of the images. We alternated two staircases with differently weighted step sizes. In the first staircase, after two consecutive correct responses the stimulus brightness was decreased by 2 steps; after 1 incorrect response the brightness was increased by 4 steps. In the second staircase, after 3 correct responses the brightness level was decreased by 3 steps, after 1 incorrect response the brightness was increased by 4 steps. Note that these are not traditional N-down/1-up procedures as the correct trial counter was not reset to zero after each pair or triplet of correct responses. However, we found in pilot work that this interleaved method stably converges to 75% correct. Brightness levels were adjusted independently for word and shape stimuli. In order to define initial brightness levels, subjects performed a 60-trial titration block for each stimulus type after the practice trials and before beginning the pre-training session. The final brightness level at the end of the titration block acted as the initial brightness level for pre-training session 1. Each subsequent session 2-10 began with the final brightness level of the previous session.

In the memory-shapes task, the number of stimuli in the encoding period was adjusted based on the average percentage correct recorded over the previous two blocks. If average performance exceeded 75% correct, one additional image was added to the encoding set. If performance dropped below 70% correct, one image was removed, down to a minimum of 2 images. We initialised the encoding set size at 4 images. Note that even though the minimum set size was 2, the underlying staircase value had no minimum value.

For the memory-words task, we employed a fixed set size of 54 words. This larger set size was based on initial pilot data and the procedure of McCurdy et al. (2013), and reflects the fact that subjects typically find encoding and remembering individual words significantly easier than encoding and remembering abstract shapes.

### Feedback

Feedback in the form of points was given based on task performance in the Control group and metacognitive calibration in the Experimental group. We rewarded the Control group on their achieved difficulty level, specified as the inverse distance between the current brightness level and the minimum brightness of 128:

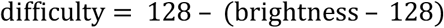

where brightness level *ϵ* [128 – 256] → difficulty level *ϵ* [0 — 128]. We chose difficulty level instead of accuracy as the relevant performance measure because accuracy was titrated to ~75% correct in each block.

We rewarded the Experimental group using the Quadratic Scoring Rule (QSR). The QSR is a proper scoring rule in the formal sense that maximum points are obtained by jointly maximizing the accuracy of choices and confidence ratings (Stael von Holstein, 1970). We mapped each confidence rating onto a subjective probability correct using a linear transformation: *p*(*correct*) = –1/3 + conf/3, where confidence rating *ϵ* [1 – 4] → *p*(*correct*) *ϵ* [0 – 1]. On each trial *i* the QSR score is then obtained as:

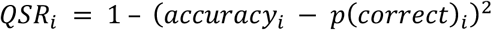

where accuracy *ϵ* [0,1] and *p*(*correct*) *ϵ* [0-1] → *QSR ϵ* [0-1]. This rule ensures that people receive the highest number of points when they are highly confident and right, or unconfident and wrong (i.e. metacognitively accurate).

Despite feedback in each group being based on different variables, we endeavoured to equate the distribution of points across groups. We used data from an initial pilot study (without feedback) to obtain distributions of expected difficulty level and QSR scores. We then calculated the average difficulty level/QSR score for each block, and fit Gaussian cumulative density functions (CDFs) to these distributions of scores. These CDFs were then used to transform a given difficulty or QSR score in the main experiment to a given number of points.

### Compensation

Participants were compensated at approximately $4 per hour, plus a possible bonus on each session. Base pay for the 60-minute pre-training session was $4, for the eight 25-minute training sessions $2 each, and for the 45-minute post-training session $3. Participants were informed they had the opportunity to earn a session bonus if they outperformed a randomly chosen other subject on that session. In practice, bonuses were distributed pseudo-randomly to ensure equivalent financial motivation irrespective of performance. All subjects received in the range of 4-7 bonuses throughout the course of the 10-session study. Bonuses comprised an additional 70% of the base payment received on any given session.

In addition to the pseudo-random bonuses, all subjects received a $3 bonus for completing half (5) of the sessions and a $6 bonus for completing all (10) of the sessions. Total earnings ranged from $37.60 – $43.90 across participants, and income did not differ significantly between groups (Control group: *mean* = $41.47; Experimental group: *mean* = $40.98; t(59) = 0.94, *p* = 0.35). The base payment was paid immediately after completing each session and accumulated bonuses were paid only if the participant completed the full 10 session experiment.

### Quantifying metacognition

Our summary measure of metacognitive calibration was the QSR score achieved by subjects before and after training. In order to separately assess effects of training on metacognitive bias (i.e. confidence level) and efficiency (i.e. the degree to which confidence discriminates between correct and incorrect trials), we also fitted meta-*d’* to the confidence rating data. The meta-*d’* model provides a bias-free method for evaluating metacognitive efficiency in a signal detection theory framework. Specifically, the ratio meta-*d’*/*d’* quantifies the degree to which confidence ratings discriminate between correct and incorrect trials while controlling for first-order performance (*d’*). Using this ratio as a measure of metacognition effectively eliminates performance and response bias confounds typically affecting other measures (Barrett et al., 2013; Fleming & Lau, 2014). We conducted statistical analyses on log(meta-*d’/d’*) as a logarithmic scale is appropriate for a ratio measure, giving equal weight to increases and decreases relative to the optimal value of meta-*d’/d’* = 1.

Meta-*d’* was fit to each subject’s confidence rating data on a per-session basis using maximum likelihood estimation as implemented in freely available MATLAB code (http://www.columbia.edu/~bsm2105/type2sdt/). Metacognitive bias was assessed as the average confidence level across a particular task and session, irrespective of correctness.

### Outline of analysis plan

By employing a combination of frequentist and Bayesian statistics, we aimed to assess the differential impact of the training manipulation across groups and the transfer of training effects across domains. In order to model the dynamics of training, we additionally assessed the drivers of the training effect using latent change score modeling and mediation analysis.

We first applied mixed-effects ANOVAs to measures of metacognition including “group” as a between-subjects factor and “task domain” as a within-subjects factor. Complementary to classical ANOVAs, we also employed a Bayesian “analysis of effects” which quantifies evidence in support of transfer of training effects across stimulus types and domains. Evidence in support of transfer is indicated by a simpler model, without stimulus or domain interaction terms, providing a better fit to the data. Finally, by modeling our data using latent changes scores, we gained insight into whether effects of training are dependent on baseline metacognitive abilities. In addition, we used mediation modelling to ask whether early shifts in confidence strategy facilitated later improvements in introspective ability.

### Analysis of effects of training

In addition to the pre-training exclusion criteria detailed above, the following set of predefined exclusion criteria was applied after data collection was complete. One subject was excluded for performing outside the range of 55 – 95% correct in at least one condition/session. One subject was excluded due to their average difficulty level calculated across all sessions dropping below 2.5 standard deviations below the group mean difficulty level. Five subjects were excluded for reporting the same confidence level on 95% of trials for 3 or more sessions. Finally, trials in which either the subject did not respond in time (response times > 2000ms) or response times were less than 200ms were omitted from further analysis (0.98% of all trials).

To evaluate effects of training, we compared data from the pre- and post-training sessions using mixed-model ANOVAs in JASP (https://jasp-stats.org/) to assess the presence of training effects as a function of domain and stimulus type (factors: [Training × Domain × Stimulus] × Group). We coded the “Stimulus” factor in terms of whether the stimulus encountered during the pre- and post-training sessions was trained or untrained. We also employed a Bayesian “analysis of effects” in JASP to quantify evidence for and against across-stimulus and across-domain transfer of training effects on confidence and metacognitive efficiency (Rouder et al., 2012).

### Latent change modeling

To assess the dependence of training gains in the (trained) perceptual domain and the (untrained) memory domain on baseline metacognitive abilities, we fit a bivariate latent change score (LCS) model to QSR scores (Kievit et al., 2017; McArdle & Nesselroade, 1994). LCS models conceptualize differences between pre- and post-training performance as latent change factors. The basic equation of the LCS model specifies the score of individual *i* in domain *Y* at post-training as a sum of the score at pre-training and a change, or difference, score:

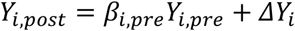

By setting the regression weight *β_i,pre_* to 1, change scores can be rewritten as follows:

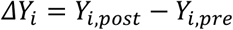

This formulation allows the change score for memory or perceptual metacognitive calibration (e.g. *ΔM* or *ΔP)* itself to be modelled as being dependent on two influences, a self-feedback process *β* and a coupling process *γ*:

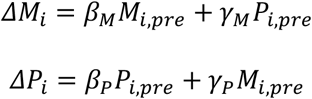

where *P* and *M* denote the QSR scores for the perceptual and memory domains, respectively. To simplify the model we included only data from the trained stimulus type in both domains. The self-feedback parameters (*β*) are assumed to reflect a combination of regression to the mean, potential dependence of training on baseline performance (e.g. the extent to which training gains are greater for individuals with low/high baseline calibration) and/or ceiling effects. The coupling parameters (*γ*) assess the extent to which change in one domain is dependent upon baseline calibration in the other domain, above and beyond the effects of self-feedback. The bivariate LCS formulation also allows estimation of the extent of correlated change, reflecting the degree to which training effects co-occur across domains, having taken into account the coupling and self-feedback parameters.

Models were estimated in the lavaan package for R (Version 5.23; Rosseel, 2012) using full information maximum likelihood, robust (Huber-White) standard errors and a scaled test statistic. We assessed overall model fit via the root-mean-square error of approximation (RMSEA; acceptable fit: < 0.08, good fit: < 0.05), the comparative fit index (CFI; acceptable fit: 0.95-0.97, good fit: > 0.97) and the standardized root-mean-square residual (SRMR; acceptable fit: 0.05-0.10, good fit: < 0.05; Schermelleh-Engel, Moosbrugger & Muller, 2003).

### Analysis of training dynamics

In order to investigate the dynamics of the training effect we calculated objective performance, metacognitive bias and metacognitive efficiency separately for each of the eight training sessions. This allowed us to visualise any progressive effects of feedback on metacognition while also establishing the stability of task performance during training sessions. To assess whether shifts in metacognitive bias mediate the impact of training on metacognitive efficiency, we fit mediation models using the Mediation Toolbox for MATLAB (https://github.com/canlab/MediationToolbox). The Mediation Toolbox uses nonparametric bootstrapping, which is more robust in handling violations to normality than traditional parametric approaches such as the Sobel test.

## Results

To quantify effects of training on both performance and metacognition, we conducted mixed-model ANOVAs comparing pre- and post-training sessions (factors: [Training × Domain × Stimulus] × Group). We coded the “Stimulus” factor in terms of whether the stimulus encountered during pre- and post-training was trained or untrained.

### First-order performance

Task performance (*d’*) was stable across pre- and post-training sessions in both groups (main effect of training: *F_1,59_* = 0.34, *P* = 0.56), and both groups performed similarly (main effect of group: *F_1. 59_* = 0.15, *P* = 0.71), as expected from the staircase procedure (Figure 2). When examining task difficulty (brightness level, controlled by the staircase procedure), we found that both groups achieved a higher difficulty level (lower brightness level) following training (main effect of training: *F_1,59_* = 15.2, *P* < 0.001), with a trend towards a more prominent difference in the Control group who received feedback on this quantity (training_control_: *F_1,31_* = 16.46, *P* < 0.001; training_experimental_: *F_1,28_* = 2.23, *P* = 0.15; training × group: *F_1,59_x* = 3.14, *P* = 0.081; Figure S1).

**Figure 2.**
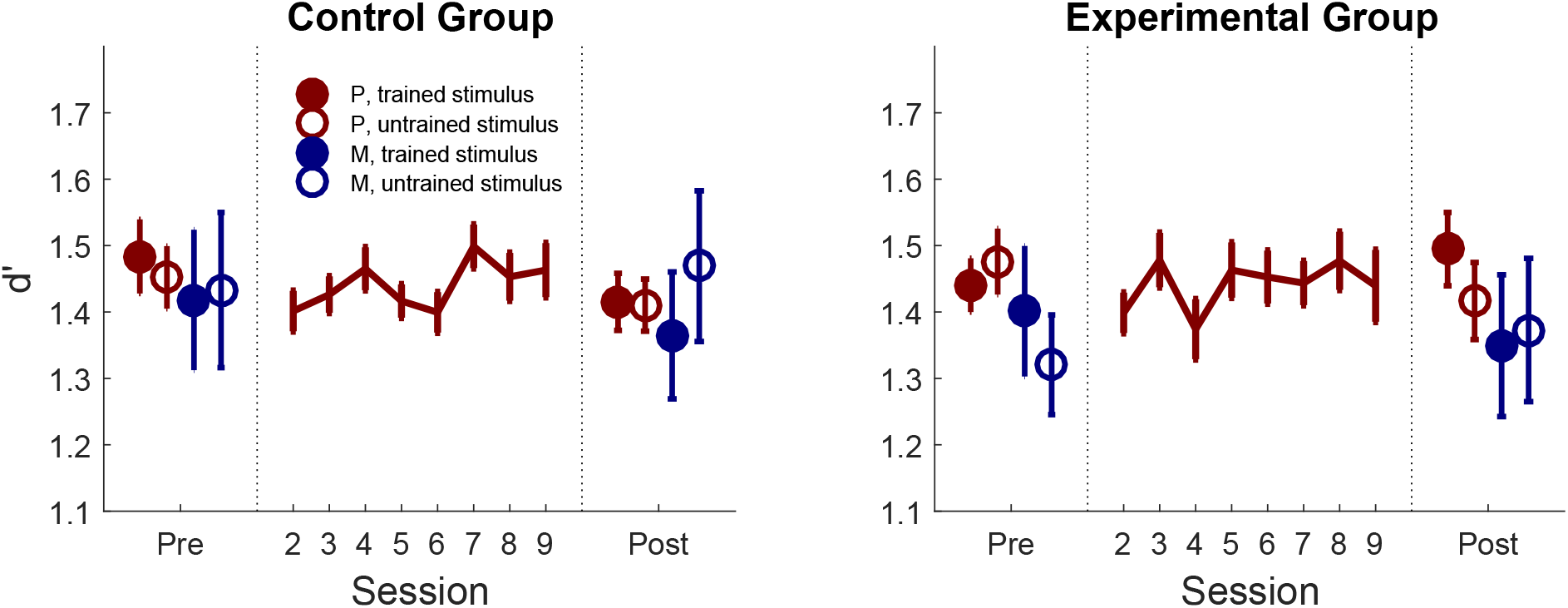
First-order discrimination performance. Effect of training on first-order performance (d’) in the control group (who received feedback on perceptual discrimination performance) and the experimental group (who received feedback on their metacognitive judgments) as a function of whether the judgment was made on a perception (red) or memory (blue) trial, and on the trained (filled) or untrained (unfilled) stimulus type. Error bars represent between-subjects SEM. P=perception; M=memory.

### Metacognitive calibration

To quantify metacognitive calibration before and after training we examined the average score achieved from the quadratic scoring rule (QSR). QSR scores are highest when confidence matches accuracy on a trial-by-trial basis – i.e. when subjects report higher confidence after correct trials, and lower confidence after errors. Critically, we observed a significant training × group interaction (*F*_1,59_ = 38.07, *P* < 0.001), driven by a robust increase in calibration in the Experimental group (*F*_1,28_ = 25.55, *P* < 0.001) and a decrease in the Control group (*F*_1,31_ = 13.15, *P* = 0.001; Figure 3 and Figure S2).

**Figure 3.**
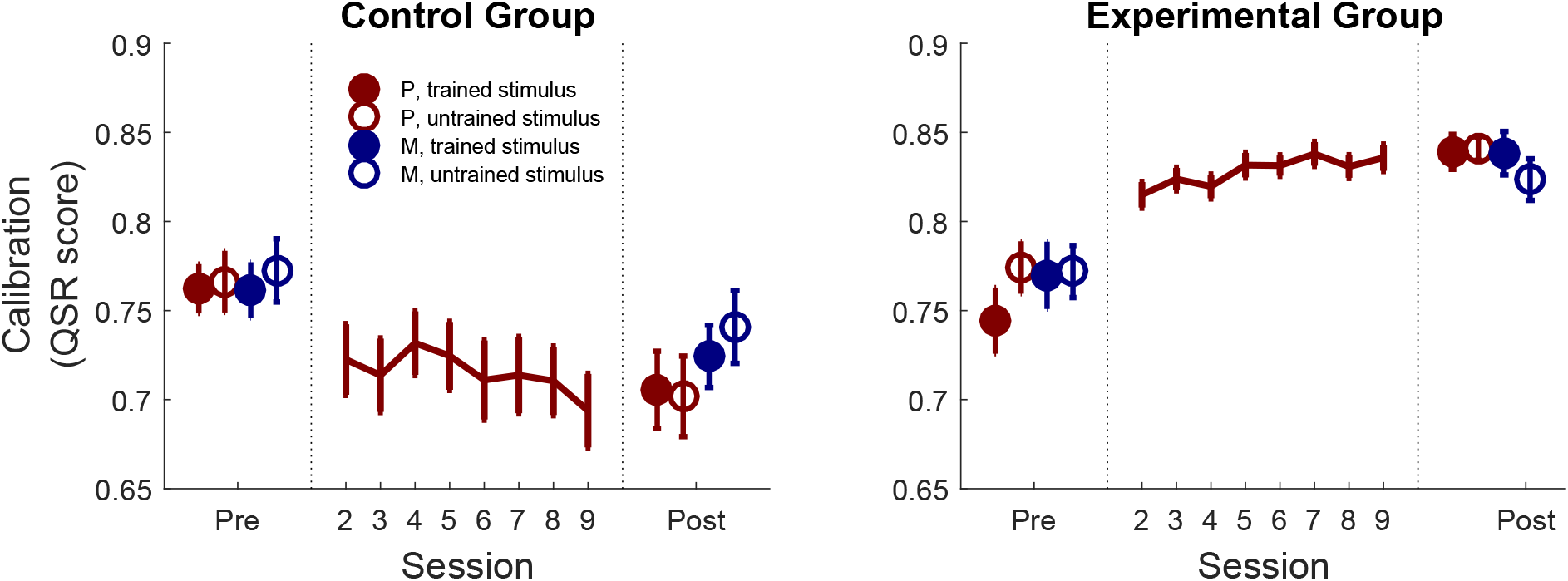
Metacognitive calibration. Effect of training on confidence calibration (the average quadratic scoring rule score, QSR). Calibration improved over training sessions in the Experimental group in the absence of changes in first-order performance (Figure 2), and this improvement transferred both to an untrained stimulus and untrained recognition memory task. Error bars represent between-subjects SEM; P=perception, M=memory.

Having revealed a selective improvement in metacognitive calibration in the Experimental group, we next asked whether this improvement generalised across stimulus types or domains. To quantify the evidence for and against across-stimulus and across-domain transfer, we performed Bayesian ANOVAs (Table 1) on QSR scores in the Experimental group. This approach (known as an “analysis of effects”; Rouder et al., 2012) analyzes all possible models of the data (e.g. main effects only, main effects + interaction effect, etc.). For each effect, a Bayes factor quantifies the degree to which the data support models including versus excluding that effect. We found evidence in support of modeling a main effect of training *(BF_inclusion_* = 1.1 × 10^10^), and evidence against modeling training × stimulus *(BF_inclusion_* = 0.13) and training × domain *(BF_inclusion_* = 0.10) interactions (Table 1, left columns). In other words, the best-fitting model is one in which the training effect on QSR scores was similar for both stimulus types (shapes and words) and both task domains (perception and memory), supporting both transfer to the untrained stimulus (within the trained perceptual task) *and* transfer to the recognition memory task, for both stimulus types. Together these results show that our metacognitive feedback protocol was able to selectively improve the correspondence between confidence and accuracy when feedback was removed, and that this improvement in confidence estimation transferred both to an untrained stimulus type and an untrained task (recognition memory).

**Table 1.**
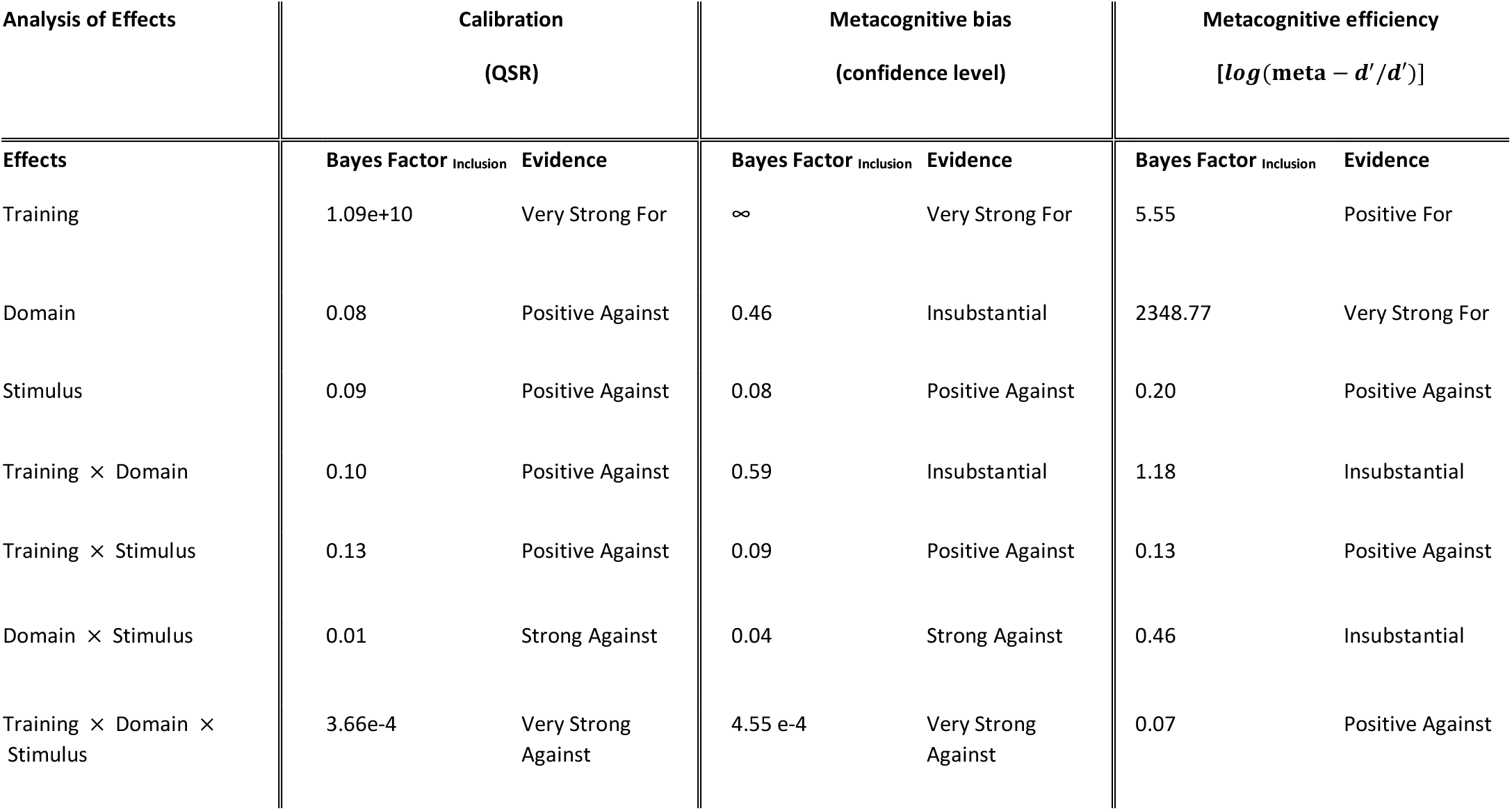
Bayesian ANOVA Analysis of Effects. Evidence in support of including different explanatory variables in models of metacognitive calibration (QSR score; left columns), metacognitive bias (confidence level; middle columns) and metacognitive efficiency (log(meta-d’/d’); right columns) in the experimental group. We obtained positive evidence against inclusion of a training × stimulus interaction term for all measures, indicating the best-fitting model is one in which the training effect is similar for both stimulus types. There was positive evidence against inclusion of a training × domain interaction term (indicating transfer across domains) in models of calibration (QSR score), and equivocal evidence for or against this term in models of both metacognitive bias and metacognitive efficiency. Strength of evidence is evaluated using Kass and Raftery’s (1995) interpretation of the Bayes Factor.

### Metacognitive efficiency and bias

Recent approaches distinguish between two key aspects of metacognitive performance (Fleming & Lau, 2014). The first is efficiency – how accurately do subjects discriminate between correct and incorrect trials for a given level of first-order task performance? The second is bias – are subjects generally more or less confident in a particular task or condition? Using a signal detection theory approach, we sought to reveal whether metacognitive improvements due to training were due to changes in efficiency, bias or both. The ratio meta-*d’*/*d’* quantifies the efficiency with which confidence ratings discriminate between correct and incorrect trials while controlling for first-order performance (*d’*) (Maniscalco & Lau, 2012). Bias was assessed as the average confidence level irrespective of whether a trial was correct or incorrect.

When analyzing metacognitive efficiency [log(meta-*d’*/*d’*)] we observed a significant training × group interaction (*F*_1,59_ = 6.96, *P* = 0.011), driven by a selective increase from pre-to post-training in the Experimental group (training_experimental_: *F*_1,28_ = 6.72, *P* = 0.015; training_control_: *F*_1,31_ = 1.39, *P* = 0.25; bottom row of Figure 4). Improvements in metacognitive efficiency were also accompanied by an overall increase in metacognitive bias (confidence level) (training_experimental_: *F*_1,28_ = 73.87, *P* < 0. 001; training_control_: *F*_1,31_ = 3.77, *P* = 0.061; training × group: F1,59 = 49.35, *P* < 0.001; top row of Figure 4).

**Figure 4.**
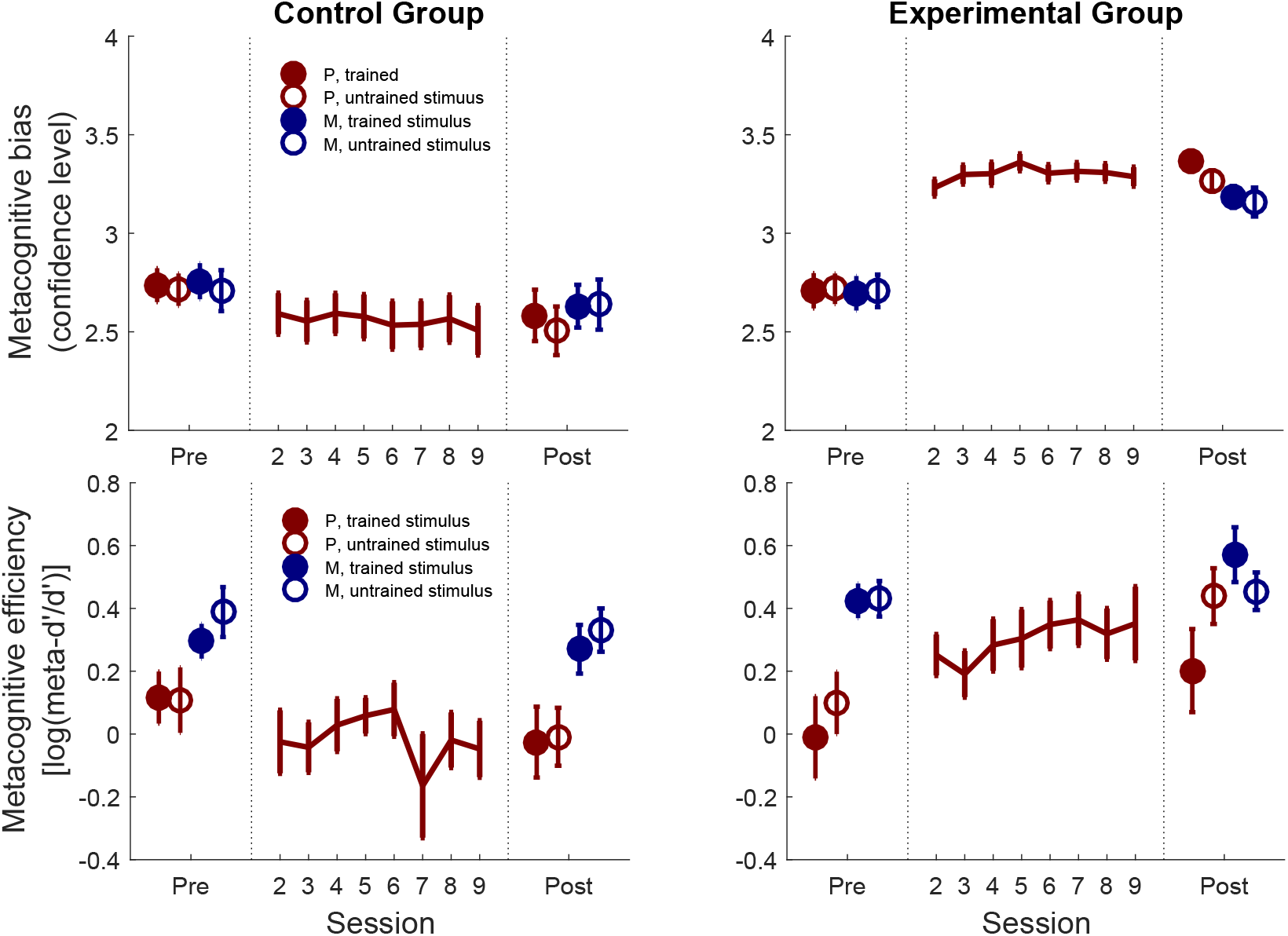
Effects of training on components of metacognition. Effects of training on metacognitive bias (confidence level; top panels) and metacognitive efficiency (log(meta-d’/d’); bottom panels). The left-hand column shows data from the Control group; the right-hand column shows data from the Experimental group. Metacognitive efficiency (log(meta-d’/d’)) gradually improved over training in the experimental group (bottom panel) in the absence of changes in first-order performance (Figure 2). Error bars represent between-subjects SEM; P=perception, M=memory. One subject was excluded when plotting mean log(meta-d’/d’) for session 6 due to a negative value of meta-d’ precluding a log-transform.

In a Bayesian analysis of effects, we found positive evidence *against* the inclusion of a training × stimulus interaction term for both metacognitive bias and metacognitive efficiency (Table 1, middle and righthand columns). In other words, the best-fitting model was one in which the training effect was similar for both stimulus types, supporting the existence of transfer to the untrained stimulus. However, there was equivocal evidence for or against transfer across domains (the training × domain interaction term) for both metacognitive bias and metacognitive efficiency, suggesting our data cannot support or refute domain-general training effects when examining these components separately.

### Latent change modeling

To identify potential drivers of improvements in metacognitive calibration we fit bivariate latent change score (BLCS) models to the QSR score data. Specifically, we examined the inter-relationship between changes in calibration for perception and memory from pre-training (T1) to post-training (T2; restricted to scores obtained for the trained stimulus type). We assessed the evidence for five possible parameters in the model. First, does baseline perceptual metacognitive ability predict the degree of change in perceptual calibration (self-feedback parameter) and/or memory calibration (coupling parameter)? Similarly, does baseline memory calibration predict the degree of change in memory calibration (self-feedback parameter) and/or perceptual calibration (coupling parameter)? Finally, is there evidence for correlated improvements (covariance of change) in perceptual and memory calibration across individuals?

Before fitting the bivariate model, we first fitted two univariate LCS models to each domain separately. In these models, the mean and variance of pre-training scores was constrained to be equal between the Experimental and Control groups. The memory model fitted the data well: *χ*^2^(2) = 0.72, *P* = 0.70; RMSEA < 0.001, 90% confidence interval [0.000, 0.265]; CFI = 1.000; SRMR = 0.083. The equivalent perceptual model revealed a poor model fit (*χ*^2^(2) = 2.43, *P* = 0.30; RMSEA = 0.084, 90% confidence interval [0.000, 0.380]; CFI = 0.91; SRMR = 0.132), which further examination indicated was driven by a higher variance of pre-training QSR scores in the Experimental compared to the Control group. Allowing the variance of T1 scores to differ between groups restored good model fit: *χ*^2^(1)= 0.62, P= 0.43; RMSEA < 0.001, 90% confidence interval [0.000, 0.439]; CFI = 1.000; SRMR = 0.046. We thus allowed perceptual T1 variance to differ between groups in the bivariate LCS model considered below. As expected, both univariate models showed evidence for positive change in QSR scores for the Experimental group (unstandardized change score intercepts – perception: 0.80, *SE* = 0.067, *z* = 11.9; memory: 0.64, *SE* = 0.086, *z* = 7.50) but not the Control group (perception: 0.19, *SE* = 0.17, *z* = 1.10; memory: 0.089, *SE* = 0.10, *z* = 0.87)^1^.

We next tested for inter-relationships between perception and memory calibration in a bivariate LCS model (shown graphically in Figure 5; significant paths are shown as thicker lines). The bivariate LCS model showed good model fit: *χ*^2^(4) = 3.20, *P* = 0.53; RMSEA < 0.001, 90% confidence interval [0.000, 0.247]; CFI = 1.000; SRMR = 0.071. Fitted model parameters are shown separately for the Control and Experimental groups in Figure 5. In addition to the expected significant latent change intercepts in the Experimental group (i.e. increasing scores), the self-feedback parameters were also positive in the Experimental group for both perception and memory, indicating that greater gains in response to training were found in individuals who started off with low metacognitive ability. Notably self-feedback effects were not observed in the Control group, indicating that this pattern of results is unlikely to be due to regression to the mean or repeated testing (constraining coupling and self-feedback parameters to be equal across groups led to a significantly worse model fit; *Δχ*^2^ (4) = 21.16, *P* < 0.001). The coupling parameter from perception at T1 to memory at T2 was also negative – individuals who started out lower in perceptual calibration improved more on memory calibration, over and above any effect of the self-feedback parameters. Finally, there was no evidence for correlated change between domains in the Experimental group. Together this analysis indicates that effects of metacognitive training depend on baseline metacognitive abilities, both within and across domains.

**Figure 5.**
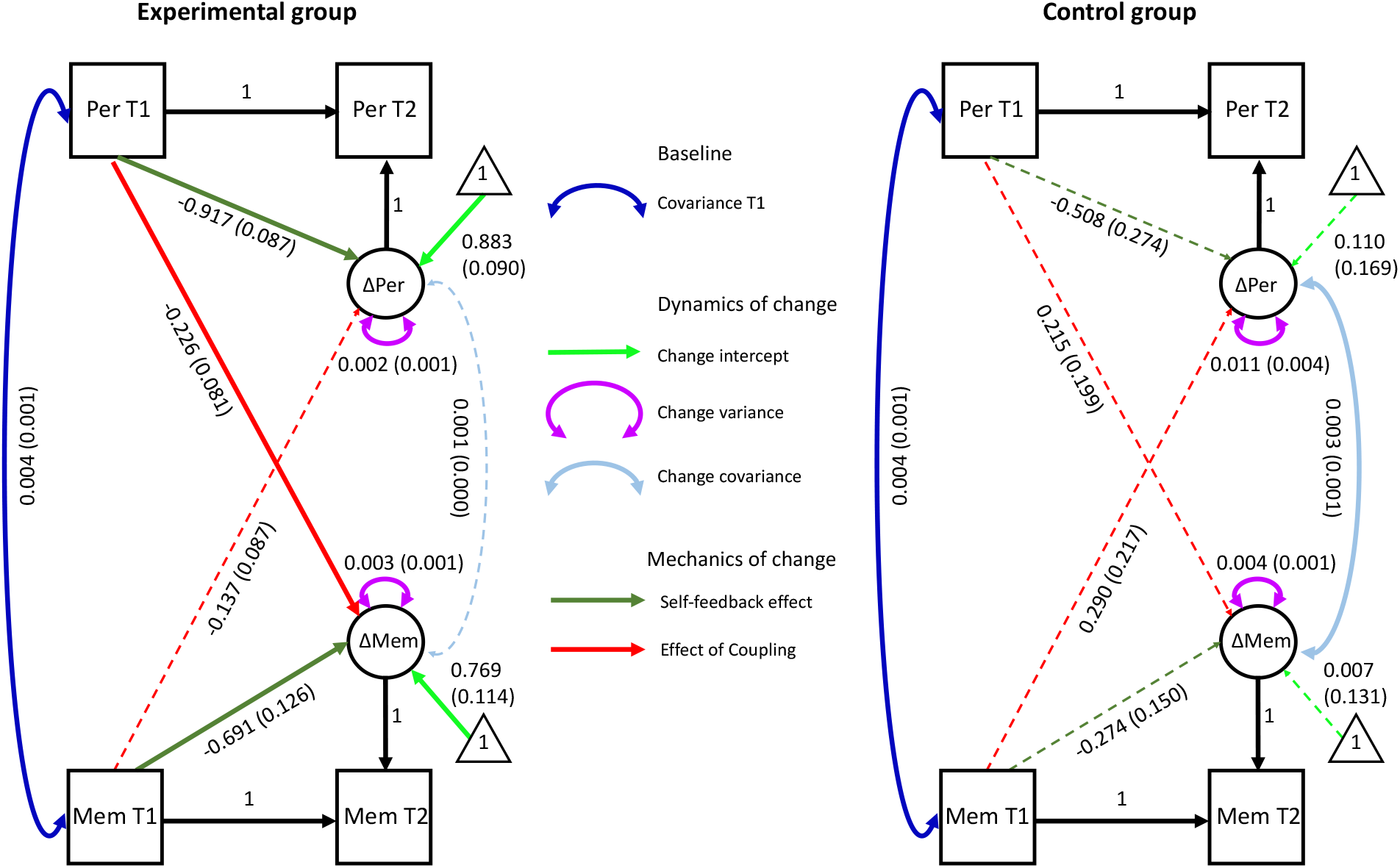
Estimated parameters for the bivariate latent change score model of metacognitive calibration (QSR scores). Calibration scores were modeled pre-(T1) and post-(T2) training across both domains, restricted to the trained stimulus type. Unstandardized parameter estimates are given separately for each group (with standard errors in parentheses). Solid lines indicate parameter significance at P < 0.05. Note that the T1 covariance, T1 intercepts and T1 memory variance were constrained to be equal across groups. T1 perception variance was estimated separately for each group as explained in the text. Per = perception; Mem = memory; T1 = pre-training; T2 = post-training.

### Dynamics of metacognitive bias and efficiency

Figure 4 indicates that a shift in metacognitive bias (confidence level) in the experimental group occurred immediately on the first training session (see also Figure S3), whereas metacognitive efficiency (meta-*d’/d’*) increased more gradually over the eight training sessions. To further quantify differences in these timecourses we calculated the session-to-session change in confidence and metacognitive efficiency (Figure 6A). The peak change in confidence was reliably earlier than the peak change in efficiency (Figure 6B; *t*(28) = 3.67, *P* = 0.001). To assess whether early changes in confidence were associated with later shifts in metacognitive efficiency, we fit a mediation model (Figure 6C). Consistent with such a hypothesis, the impact of feedback type (i.e. group) on increases in log(meta-*d’/d’*) was positively mediated by initial shifts in confidence (*t*(58) = 2.24, *P* = 0.028).

**Figure 6.**
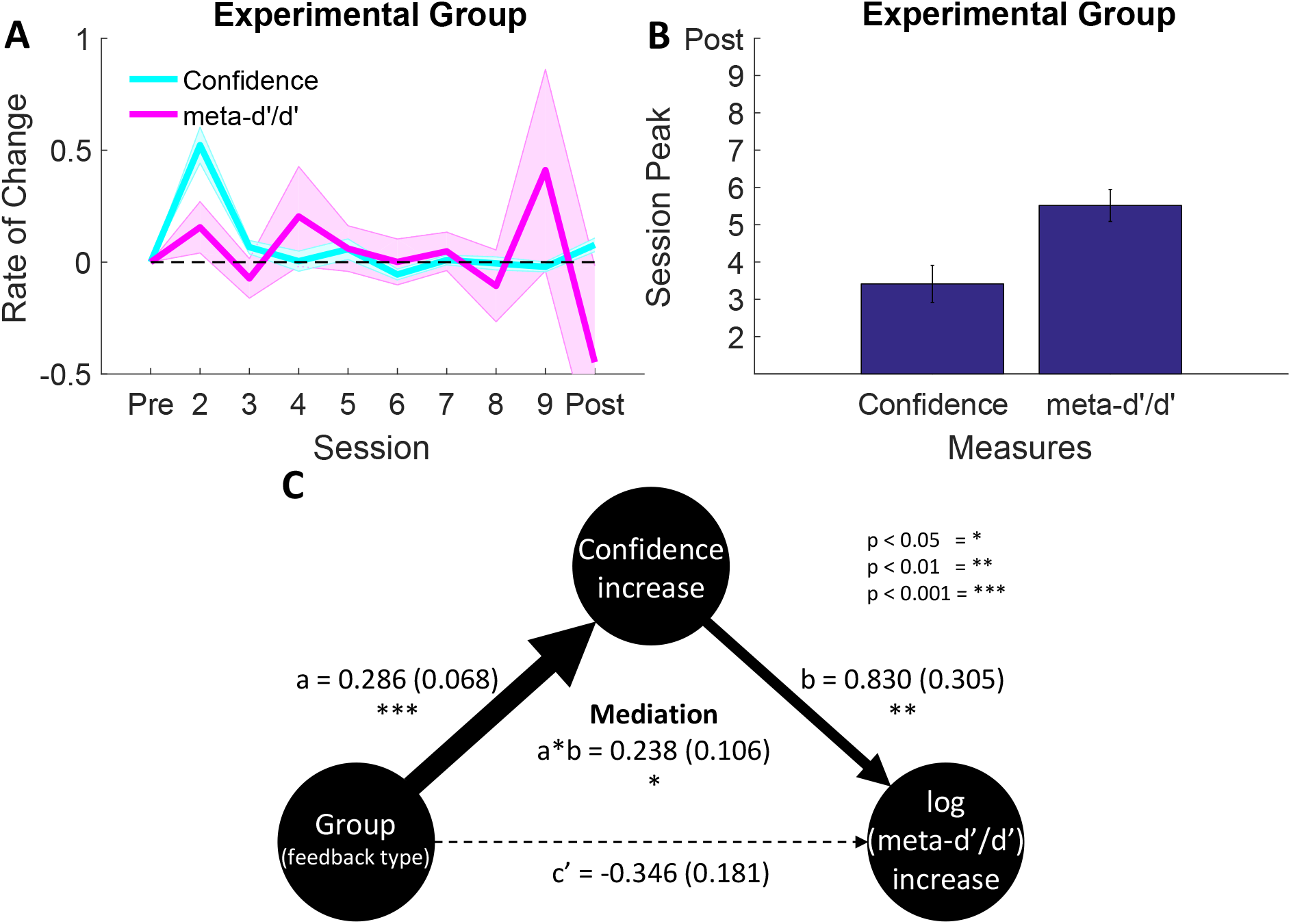
Temporal dissociation of shifts in metacognitive bias and metacognitive efficiency. **(A)** Rate of change over sessions of confidence level and meta-d’/d’ in the experimental group showing an early shift towards responding with higher confidence (see also Figure S3). This shift in confidence was dissociated in time from a more gradual improvement in metacognitive efficiency, with the largest changes occurring towards the end of training. **(B)** The session at which this peak shift occurred was significantly earlier for metacognitive bias (confidence level) compared to metacognitive efficiency (meta-d’/d’). **(C)** Early increases in confidence mediate the impact of feedback type on later increases in metacognitive efficiency. Values outside of parentheses indicate the coefficient mean and values inside parentheses indicate the SEM.

## Discussion

Here we reveal a domain-general enhancement of metacognitive abilities despite objective performance (*d’*) remaining unchanged across two distinct perceptual and memory tasks. These changes were only observed when feedback was targeted to metacognitive judgments – an active control group who performed the same tasks but received feedback on first-order (objective) performance did not show the same improvement. Since feedback and financial incentives were matched across groups, motivational factors are unlikely to account for our results. Our findings are instead consistent with a specific effect of metacognitive feedback in enhancing subjects’ ability to introspect about self-performance.

In addition to a main effect of training on a trained stimulus type, we obtained evidence that improvements in calibration scores generalized both to other instances of brightness discrimination and, more importantly, an untrained task (recognition memory). This result indicates that the feedback individuals receive on their confidence-accuracy relationship on one task can lead to improved confidence calibration for unrelated tasks, after feedback is removed. Current evidence for a shared neurocognitive resource for metacognition is ambiguous, partly due to a difficulty of distilling metacognitive processes from those supporting primary task performance (Ais et al., 2016; Baird et al., 2013; Song et al., 2011; McCurdy et al., 2013). The observation of domain-general enhancement provides a novel perspective on this issue, suggesting the existence of generic metacognitive resources that can be altered through training. Previous work has suggested confidence estimates are compared in a “common currency” across a range of decision scenarios (de Gardelle & Mamassian, 2014; Faivre et al., 2017), and training may boost the fidelity of such shared signals. In turn our findings hold promise for the future development of training protocols to boost metacognition in applied settings, in which administering domain-specific adaptive training protocols may facilitate improvements in metacognitive abilities more generally.

Latent change score modeling of QSR scores indicated that baseline performance in both trained and untrained tasks (perception and memory) predicted the extent of training gains, with lower baseline levels in a particular domain predicting greater training gains in that domain. In addition, there was evidence for a cross-domain coupling in which lower initial scores on the trained (perceptual) task predicted greater gains in the untrained memory task, over and above effects of self-feedback. These effects were not observed in the active control group, making explanations of such dynamics in terms of regression to the mean or repeated practice less likely. Interestingly a similar pattern has been observed in the literature on working memory training, with the largest training gains observed for those initially low in WM capacity (Zinke et al., 2012; 2014; although see Bissig & Lustig, 2007). Such findings are potentially consistent with initially low performing individuals having a larger (underused) latent potential for WM/metacognition, therefore leading to a stronger response to training. A less interesting explanation is that there are ceiling effects on potential QSR scores, leading to a natural slowdown in gains as a function of starting point. Future work (for instance examining the effects of training over multiple time points, and/or with larger *N* to more precisely estimate the dynamics and cover a wider range of ability levels) is needed to disentangle these possibilities.

We also examined how two key components of metacognition – metacognitive efficiency (meta-*d’/d’*) and metacognitive bias (confidence level) – evolved over the course of training. For both components, we observed significant effects of training in the experimental group. However, when examining transfer for each component individually, the picture was more mixed than for the composite calibration measure: while both components generalised to other instances of brightness discrimination, there was equivocal evidence for across-domain transfer to memory metacognition. This pattern of results is potentially consistent with a domain-specificity of metacognitive efficiency for perception vs. memory (McCurdy et al., 2013; Baird et al., 2013; Fleming et al., 2014), and recent observations that metacognitive efficiency, while stable within a particular subject across sessions, may be idiosyncratic to particular tasks (Ais et al., 2016). However, we note initial metacognitive efficiency scores for the memory task were high, potentially leading to a ceiling effect on subsequent improvement in this domain. In addition, it remains to be determined whether enhancements of perceptual metacognitive efficiency are limited in transfer to other features within the same modality (such as visual contrast and orientation; Song et al., 2011) or also generalise to other perceptual modalities, such as audition (Faivre et al., 2017).

The timecourse of training effects provides insight into potential mechanisms supporting metacognitive improvement. While confidence levels increased during the very first training session and remained stable throughout the remainder of the experiment, metacognitive efficiency climbed more gradually across the eight training sessions. One possible account of this pattern (supported by a mediation analysis) is that an initial shift in confidence strategy facilitates later increases in metacognitive efficiency allowing, for instance, higher confidence to be effectively targeted to correct trials (Figure S2). An implicit signal of whether a first-order decision is likely to be correct may then gradually become associated with higher confidence reports over time, and reinforced by the feedback schedule.

It is important to note that an initial shift in confidence bias does not necessarily reflect a change in metacognition, and may instead reflect a strategic shift in response to the onset of feedback protocol and instructions. Critically, however, such a strategic shift alone is unlikely to explain later change in metacognitive efficiency. To establish the expected impact of a non-specific bias on measures of metacognitive efficiency, we conducted numerical simulations in which the pre-training confidence data were shifted to create an artificial bias in confidence level (Figure S6). These simulations show that “learning” to increase mean confidence leads to an increase in calibration score, as expected, but is insufficient to produce the observed increases in metacognitive efficiency. Indeed, when confidence bias is artificially induced, metacognitive efficiency is expected to be lower post-compared to pre-training – precisely the opposite of what we find. Thus we believe that these simulations lend support to a conclusion that metacognitive efficiency is specifically increased following feedback on metacognitive judgments, and this effect is not a trivial consequence of strategic biases in confidence.

Our work goes significantly beyond previous attempts to improve the resolution or calibration of confidence judgments. Adams and Adams (1958), Lichtenstein et al. (1982), and Sharp et al. (1988) all reported changes in the confidence-accuracy relationship for participants who received feedback on the correctness of their confidence ratings but lacked active control groups or controls for changes in performance (although Sharp et al., 1988, were aware of this issue). Indeed, participants in the feedback condition of Adams & Adams (1958) reported feeling markedly more enthusiastic about the experiment, suggesting motivation differences may have confounded effects of feedback. Here we addressed this concern by matching feedback schedules and first-order performance levels between the experimental group and an active control group, who received equivalent feedback directed at first-order performance. Intriguingly, the feedback protocol implemented in the present study may represent one among many possible methods for inducing increases in metacognitive efficiency. Other feedback protocols may operate via a different mechanism, e.g. learning to decrease error trial confidence, rather than increasing one’s confidence in being correct. Future work could investigate the scope of possible training protocols by manipulating parameters such as titrated performance level and feedback schedule.

Fine-grained introspective ability is useful for several reasons. First, it aids the control of task performance – becoming aware of making suboptimal choices is a useful signal for prompting changes of mind (Folke et al., 2017) and for the guidance of learning (Metcalfe & Finn, 2008; Nietfeld & Schraw, 2002; Purcell & Kiani, 2016). Second, appropriate sensitivity to self-performance is important when interacting with others (Bahrami et al., 2010; Shea et al., 2014), allowing communication of degrees of belief to improve group decision-making and avoid overconfident testimony (e.g. in an eyewitness context; Busey et al., 2000). Finally, metacognition is a potential target of interventions in psychiatric disorders including schizophrenia and depression (Moritz & Woodward, 2007). Developing tools to improve metacognitive abilities may therefore have widespread impact in a variety of settings. Here, despite obtaining evidence for generalization to an untrained task, such “transfer” was limited to a suite of computerized, 2-alternative forced choice tasks with confidence ratings. Further work is needed to assess whether metacognitive training has more widespread benefits for unrelated tasks and/or for learning contexts that place demands on metacognitive control.

Our results open up new questions regarding the nature of the malleability of metacognition displayed in the present study. Specifically, the duration and generality of improvements in introspective abilities remain to be determined. We might expect improvements in the ability to introspect about self-performance to be accompanied by changes in brain structure, function, and/or connectivity within frontoparietal networks previously implicated in supporting metacognition (Fleming et al., 2010; Fleming and Dolan, 2012; McCurdy et al., 2013; Baird et al., 2013; Allen et al., 2016; Cortese et al., 2017). A distinction has recently been drawn between lower-level (and potentially generic) signals of confidence and higher-order elaboration of such signals for use in communication and control (Fleming & Dolan, 2012; Morales et al., 2018). By combining the current behavioural intervention with neuroimaging measures it may be possible to determine whether one or both of these levels of processing are affected by metacognitive training. Ongoing work in our laboratory is tackling this question.

## Author contributions

J.C., H.L. and S.M.F. developed the study concept. All authors contributed to the study design. Task programming and data collection was conducted by J.C. J.C. performed data analysis and interpretation under supervision of S.M.F. S.M.F. and R.A.K. carried out latent change score modeling. J.C. and S.M.F. drafted the manuscript, and M.T.S., A.K.S., R.A.K. and H.L. provided critical revisions. All authors approved the final version of the manuscript for submission.

## Acknowledgements

These results were previously disseminated as a poster presentation at the Society for Neuroscience Annual Meeting (2017). This work was supported by the National Institute of Neurological Disorders and Stroke of the National Institutes of Health (Grant No. R01NS088628) to H.L. and S.M.F. S.M.F. is supported by a Sir Henry Dale Fellowship jointly funded by the Wellcome Trust and Royal Society (206648/Z/17/Z). The Wellcome Centre for Human Neuroimaging is supported by core funding from the Wellcome Trust (203147/Z/16/Z). A.K.S and M.T.S. are grateful to the Dr. Mortimer and Theresa Sackler Foundation, which supports the Sackler Centre for Consciousness Science. Anonymised behavioural data and code for reproducing all analyses in the manuscript can be obtained at https://github.com/metacoglab/CarpenterMetaTraining.

## Footnotes

1 Note that these intercept parameters can be interpreted only in the context of the full LCS model that includes the self-feedback pathway.

